# Neuroticism is linked to cognitive decline and increased risk of Alzheimer’s disease through dysregulation of excitatory neurons

**DOI:** 10.64898/2026.06.27.735015

**Authors:** Sneha Sharma, Natacha Comandante-Lou, Yiyi Ma, Masashi Fujita, David A. Bennett, Andrea R. Zammit, Philip L. De Jager

## Abstract

**INTRODUCTION:** Neuroticism is an established risk factor for Alzheimer’s disease (AD), yet the molecular mechanisms linking this personality trait to neurodegeneration remain poorly understood.

**METHODS:** We leveraged single-nucleus RNA-sequencing data from longitudinal cohort studies of cognitive aging (n = 655) to investigate the mechanisms mediating the association between neuroticism and AD.

**RESULTS:** We identified two genes associated with neuroticism, both of which are downregulated in excitatory neurons: *ADRA1B* and *LY6E-DT*. In addition, we found that neuroticism is associated with enrichment of a specific excitatory neuron subpopulation: Exc.12. Further analysis revealed that Exc.12 partially mediates the relationship between neuroticism and AD, accounting for 12.2% of the total effect.

**DISCUSSION:** The dysregulation of excitatory neurons may represent a key cellular change that links neuroticism to AD.

## Introduction

Neuroticism is a personality trait characterized as a heightened sensitivity to stress. It remains relatively stable across adulthood and reflects enduring differences in vulnerability to chronic stress^1^. Previous studies have shown that individuals with higher levels of neuroticism are at increased risk of developing Alzheimer’s disease (AD) later in life^2^. Large prospective cohort studies have demonstrated that this relationship persists across decades of follow-up, suggesting that neuroticism may contribute to disease development rather than simply reflecting a prodromal symptom of emerging dementia^3,4^.

Despite the established association between neuroticism and AD, major questions remain regarding the molecular mechanisms underlying this relationship. In our prior work, we used bulk RNA-sequencing data to assess the effect of personality on the brain transcriptome and found that neuroticism was associated with a set of gene modules, supporting a role for transcriptional changes as mediators of neuroticism’s effects^5^. While these findings provided initial insights into the transcriptomic impact of neuroticism, they were limited by the use of bulk tissue and a focus on broad gene modules, which obscured cell-type-specific and individual gene-level associations.

In this study, we expand on our previous work by investigating the relationship between neuroticism and AD within neocortical cell types, leveraging single-nucleus RNA-sequencing (snRNA-seq) data from 655 individuals enrolled in a longitudinal cohort study of cognitive aging^6^. Using these data, we first reproduced the association between neuroticism and AD in this cohort. Then, we performed differential expression analyses to capture gene expression changes associated with neuroticism within cell types. Informed by these cell-type-specific findings, we prioritized and tested potential mediators of the relationship between neuroticism and AD, identifying a specific cell subpopulation of interest. By examining AD through the lens of neuroticism, this study elucidates mechanisms linking this personality trait to neurodegeneration and highlights a cellular target for further investigation.

## Materials and methods

### Study participants

Data were obtained from two longitudinal cohort studies of cognitive aging with prospective brain donation: the Religious Orders Study and the Rush Memory and Aging Project, referred to jointly as ROSMAP^6^. The two studies were designed to be analyzed together and are led by the same team of investigators, using shared ante-and post-mortem phenotypic characterization protocols at the item level. Participants did not have a diagnosis of dementia at enrollment and underwent detailed cognitive testing annually until death. Participants signed an informed consent, a repository consent, and an Anatomical Gift Act to donate their brains after death. Both studies were approved by an Institutional Review Board at Rush University Medical Center. The subset of ROSMAP participants included in this study had been evaluated for neuroticism using the NEO Five-Factor inventory and had been profiled using snRNA-seq, culminating in a sample size of 655 participants.

### Phenotype assessment

Neuroticism was assessed at baseline using 12 items from the NEO Five-Factor Inventory. Each participant completed an interviewer administered survey, rating agreement with each item on a 5-point Likert scale. Participants received a cumulative neuroticism score ranging from 0 to 48, with higher scores indicating greater neuroticism. Scores were also calculated at the facet-level, with each participant receiving a score for anxiety, anger/hostility, depression, self-consciousness, impulsiveness, and vulnerability to stress^2^.

Global cognition was assessed by averaging participant z-scores from a range of cognitive tests capturing episodic, semantic, and working memory, as well as visuospatial ability and perceptual speed^7^. Cognitive change was ascertained by calculating a person-specific rate of change in global cognition over time; the rate of cognitive change was estimated from a linear mixed-effects model using global cognition as the longitudinal outcome and adjusting for age at baseline, sex, and years of education.

Clinical assessment of cognitive health was conducted each year, blinded to prior years. At the time of death, a neurologist with expertise in dementia reviewed pertinent clinical data and rendered a summary diagnostic opinion regarding the most likely cognitive diagnosis: no cognitive impairment, mild cognitive impairment, Alzheimer’s dementia, or other primary cause of dementia. This diagnostic impression was made blinded to neuroticism data and neuropathology data^8^.

### Neuropathologic assessment

After death, a structured neuropathologic examination was performed to measure Aβ and tau pathology in the brain^9^. Aβ pathology was identified by immunohistochemistry and quantified using image analysis to calculate the percentage area of the cortex occupied by Aβ protein. Tau pathology was identified by immunohistochemistry using the AT8 antibody and cortical density per mm^2^ was determined by systematic sampling of the cortex. Aβ and tau pathology data were square-root transformed to fit a normal distribution.

### Transcriptomic data preparation

After death, samples from the dorsolateral prefrontal cortices of ROSMAP participants were obtained for snRNA-seq. The methods by which the snRNA-seq data were generated, quality controlled, and pre-processed are described in detail in prior reports^10–12^. Briefly, nuclei were isolated from pooled grey matter samples and processed via the 10x Chromium platform. Sequencing reads were aligned using CellRanger^13^ and noise was removed using CellBender^14^, followed by demultiplexing via demuxlet^15^ to assign nuclei to individual participants based on genotype data. The RNA count matrices underwent normalization, scaling, and dimensionality reduction using Seurat^16^. Cell types were annotated using a weighted ElasticNet-regularized logistic regression classifier trained on a pre-existing atlas^17^. Quality control involved applying cell-type-specific UMI and gene thresholds and using a multi-method approach to identify and remove doublets.

The samples were sequenced at differing time points and institutions, culminating in three datasets we refer to as CUIMC1, CUIMC2, and MIT. We harmonized the three snRNA-seq datasets to ensure that only one instance of each participant exists in the overall sample. Cells from all three datasets were classified using the same cell atlas, allowing for integrated analysis.

### Phenotype-based Regression Analysis

Logistic regression modeling was performed to assess the association between neuroticism and AD diagnosis at death. Linear regression modeling was conducted to quantify the association between neuroticism and cognitive change. Similarly, linear regression modeling was conducted to evaluate the association between neuroticism and Aβ as well as tau pathology. Confounders including sex, years of education, age at death, and study (ROS vs. MAP) were adjusted for in the models. The summary statistics of the regression analyses are provided in **Supplementary Table S1**.

### Differential Expression Analysis

To identify genes associated with neuroticism for specific neocortical cell types, a differential expression analysis was performed for each dataset. The input for the differential expression analysis was a pseudobulk expression matrix, generated by aggregating UMI counts for each gene across cells in each cell type. The cell types used for pseudobulking included: astrocytes, microglia, inhibitory neurons, excitatory neurons, oligodendrocytes, oligodendrocyte precursor cells, and vascular niche cells. The pseudobulk matrix was filtered for subjects with at least 10 cells and for genes expressed in at least 10% of subjects.

To account for the mean-variance relationship across genes, the *voom* function^18^ from the limma package was applied to generate a set of precision weights. These weights were then used to fit a linear model for each gene with the *lmFit* function^18^, adjusting for age at death, sex, years of education, study, and post-mortem interval. The output statistics of the linear model were computed using the *eBayes* function^18^.

Employing a two-stage design, genes reaching nominal significance (p <= 0.05) in the Stage 1 dataset (CUIMC1) were advanced to meta-analysis. A fixed-effects meta-analysis of the Stage 2 datasets (CUIMC2 and MIT) was performed using the *rma* function from the metafor package^19^. Correction for multiple hypothesis testing was done on resulting p-values from the fixed-effects meta-analysis using the Benjamini-Hochberg adjustment method. Genes with an adjusted p-value <= 0.05 were considered to have a statistically significant association with neuroticism. The summary statistics for the differential expression analysis are available in **Supplementary Table S2 & S3**.

### Cell Subpopulation Analysis

Cell subpopulations within cell types were annotated using a reference from a published cell atlas of the aged prefrontal cortex^10^. Cell subpopulations were labeled using the Seurat query mapping pipeline. Details regarding the subpopulation mapping methodology are provided in a previous report^12^.

To assess differential gene expression at the subpopulation level, pseudobulk matrices were generated by aggregating UMI counts for each gene across all cells of that subpopulation. Each pseudobulk matrix was filtered for subjects with at least 10 cells and for genes expressed in at least 10% of subjects. The R-based pipeline detailed in the section above was employed to identify differentially expressed genes in each subpopulation, for each dataset. A fixed-effects meta-analysis of all three datasets (CUIMC1, CUIMC2, and MIT) was performed to generate the summary statistics for each gene, within each subpopulation. The summary statistics for the differential expression analysis at the subpopulation level, filtered for genes of interest, are available in **Supplementary Table S4**.

Cell subpopulation frequencies were calculated within each cell type. We define cell subpopulation frequency as the number of cells of the subpopulation of interest (ex: Exc.12) divided by the total number of cells of the respective cell type (ex: excitatory neurons), multiplied by 100 to yield a percentage value.

To assess cell type composition shifts in response to gene expression changes, the association between expression levels of neuroticism-associated genes vs. cell subpopulation frequency was examined using linear regression, adjusting for demographic confounders (age at death, education, sex, study), as well as technical confounders (post-mortem interval, cell count, median genes detected, and percentage of mitochondrial genes). The output of these regression analyses is available in **Supplementary Table S5**.

Additionally, the associations between cell subpopulation frequency vs. neuroticism score, AD diagnosis at death, cognitive decline, Aβ and tau pathology were examined using regression modeling, adjusting for the demographic and technical confounders listed above. The output of these analyses is available in **Supplementary Tables S6 and S7**.

### Mediation analysis

Mediation modeling was performed to assess whether a specific cell subpopulation of interest mediates the relationship between neuroticism and AD diagnosis at death. The *mediate* function^20^ was run with bootstrapping for 30000 simulations to estimate casual mediation effects. Mediation modeling was also used to assess whether the cell subpopulation of interest mediates the relationship between neuroticism and cognitive change, as well as the relationship between neuroticism and AD pathology. The results of the mediation analyses are available in **Supplementary Table S8**.

## Results

### Neuroticism is reproducibly associated with AD and cognitive decline

First, we used logistic regression to reproduce the relationship between neuroticism and AD in our sample of 655 ROSMAP participants (**Table 1**). We confirmed that neuroticism is positively associated with a clinical diagnosis of AD (*p* = 3.69 × 10^−4^), adjusting for sex, years of education, age at death, and study. Notably, this association remained significant after controlling for Aβ pathology (*p* = 9.36 × 10^−4^) and tau pathology (*p* = 3.06 × 10^−3^). Additionally, we validated that neuroticism is associated with an increased prevalence of AD dementia at death (*p* = 8.02 × 10^−5^) (**Fig. 1A**).

**Figure 1:**
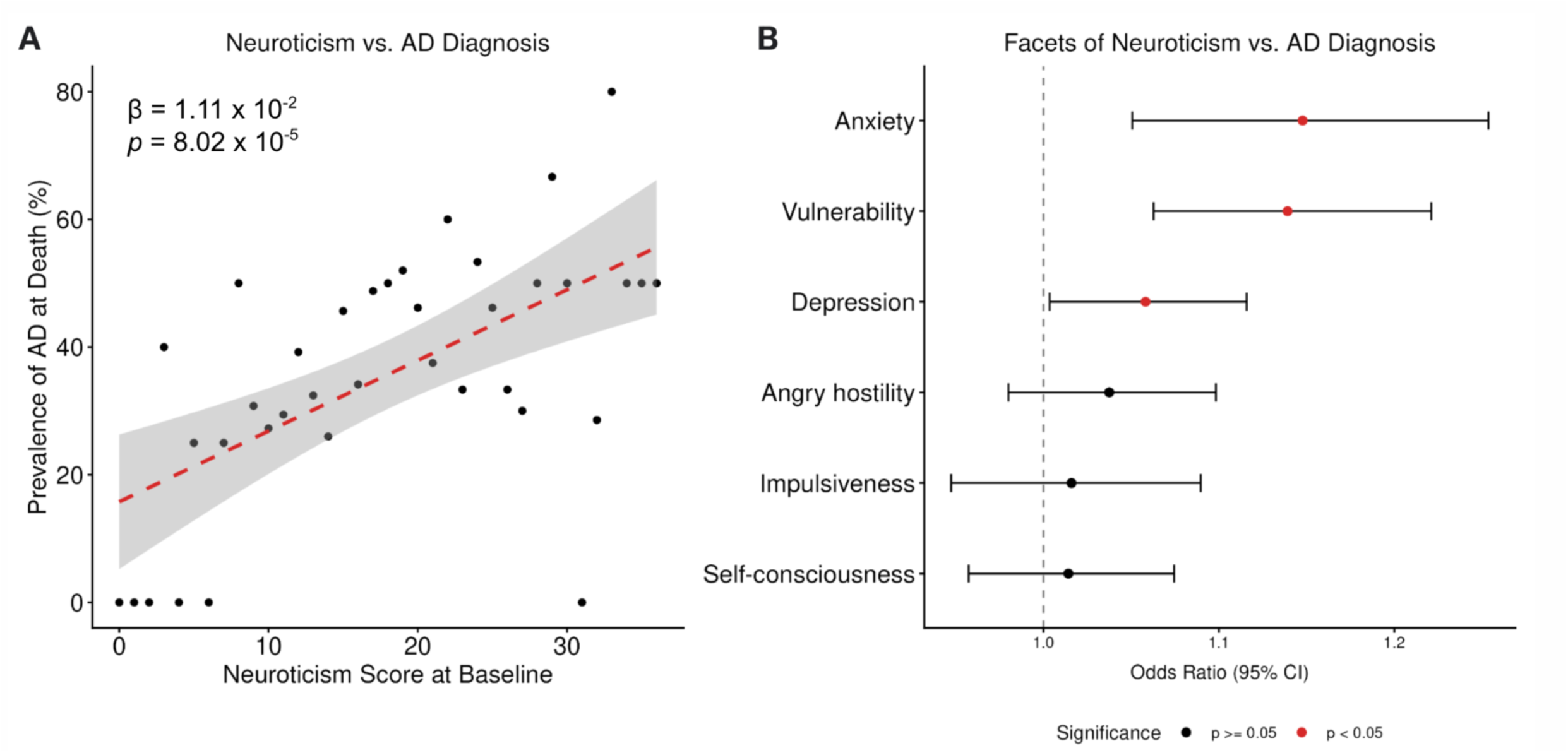
Neuroticism is significantly and reproducibly associated with AD in our sample. (A) A scatter plot illustrating neuroticism score at study enrollment vs. the percentage of patients with an AD diagnosis at death. To calculate the percentage shown on the y-axis, participants were grouped by neuroticism score; the number of individuals with an AD diagnosis at death was then divided by the total number of participants in that group. The red dashed line represents a linear regression model fit to the data; the shaded region represents the 95% confidence interval, indicating the precision of the estimated effect across the range of neuroticism scores. (B) A forest plot visualizing the relationship between individual facets of neuroticism vs. AD diagnosis at death. Each point represents an odds ratio, with red points denoting statistical significance. The error bars indicate the 95% confidence interval. The associations between facets of neuroticism vs. AD diagnosis were quantified using logistic regression modeling, adjusting for age at death, years of education, and sex.

**Table 1:** Phenotype and demographic characteristics of ROSMAP participants included in analysis, stratified by sequencing cohort.

| **Characteristic** | **CUIMC1**<br>N = 371<sup>1</sup> | **CUIMC2**<br>N = 172<sup>1</sup> | **MIT**<br>N = 112<sup>1</sup> |
| --- | --- | --- | --- |
| **Neuroticism Score** | 16 (7) | 17 (6) | 17 (6) |
| **Cognitive Diagnosis** |  |  |  |
| No/Mild CI | 225 (61%) | 94 (55%) | 73 (65%) |
| AD | 135 (36%) | 76 (44%) | 39 (35%) |
| Non-AD Dementia | 11 (3.0%) | 2 (1.2%) | 0 (0%) |
| **TAU pathology**<sup>2</sup> | 2.12 (1.30) | 2.28 (1.17) | 1.92 (1.45) |
| **Aβ pathology**<sup>2</sup> | 1.70 (1.19) | 1.81 (1.05) | 1.29 (1.22) |
| **Race** |  |  |  |
| Black | 0 (0%) | 0 (0%) | 2 (1.8%) |
| White | 371 (100%) | 172 (100%) | 110 (98%) |
| **Sex** |  |  |  |
| Female | 255 (69%) | 105 (61%) | 56 (50%) |
| Male | 116 (31%) | 67 (39%) | 56 (50%) |
| **Education (years)** | 16.6 (3.4) | 16.9 (3.6) | 17.5 (3.6) |
| **Age at Death (years)** | 89 (7) | 90 (7) | 86 (6) |
| **PMI (hours)** | 7.7 (5.2) | 8.4 (5.5) | 8.3 (8.9) |
| <sup>1</sup> Mean (SD); n (%) |  |  |  |
| <sup>2</sup> Square-root transformed |  |  |  |

We then investigated whether specific facets of neuroticism drive the overall association with AD. Our facet-level analysis revealed that vulnerability to stress (*p* = 2.37 × 10^−4^), anxiety (*p* = 2.24 × 10^−3^), and depression (*p* = 3.68 × 10^−2^) are positively associated with a clinical diagnosis of AD, whereas angry hostility, impulsiveness, and self-consciousness are not significant contributors (p > 0.05) to the overall neuroticism-AD association (**Fig. 1B**).

Furthermore, we validated that neuroticism was significantly associated with the slope of cognitive decline over time (*p* = 2.91 × 10^−4^) (**Fig. 2A**). Of the core cognitive domains, semantic memory (*p* = 1.16 × 10^−3^) and episodic memory (*p* = 9.00 × 10^−4^) were most significantly impacted by neuroticism (**Fig. 2B**).

**Figure 2:**
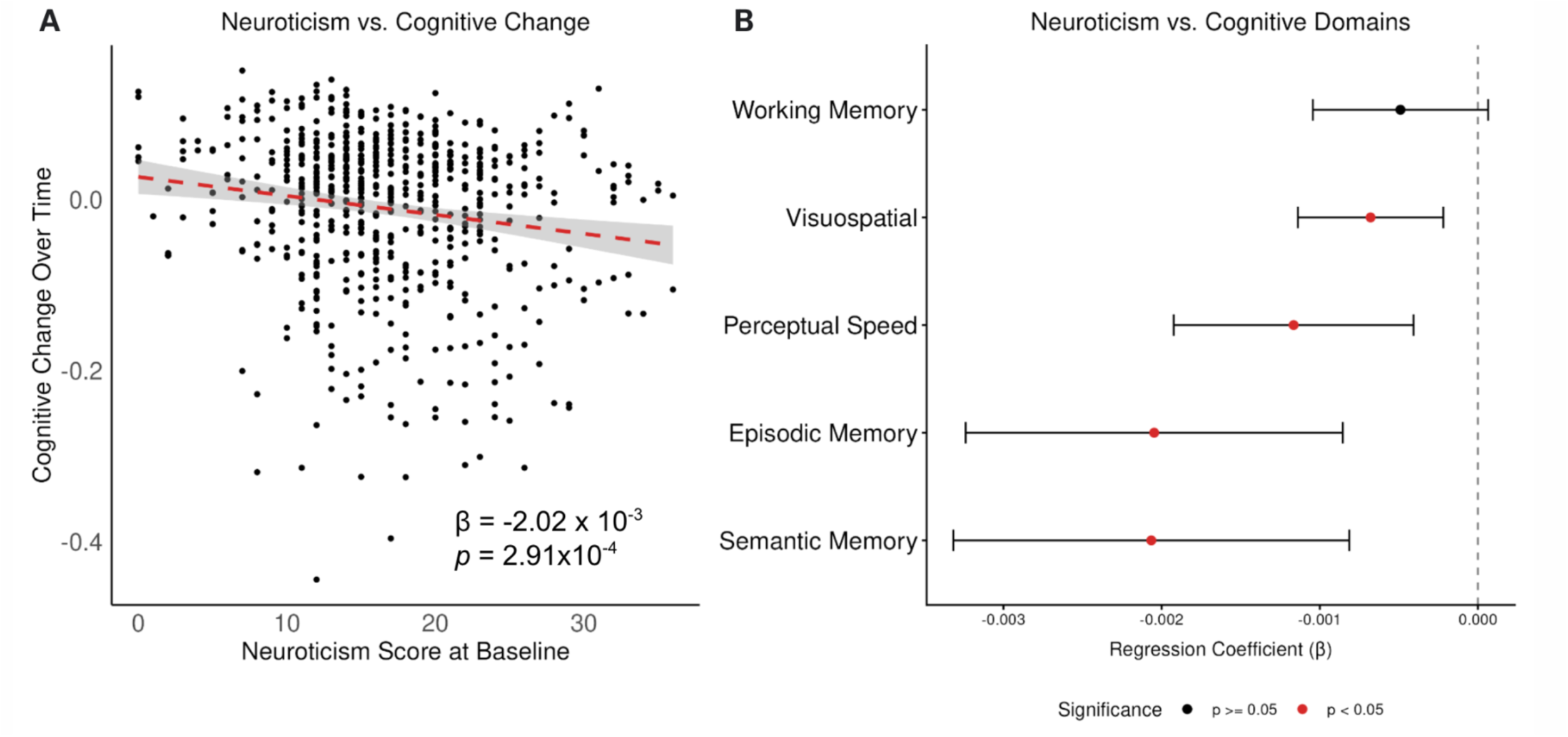
Neuroticism significantly impacts key cognitive domains of memory. (A) A scatter plot illustrating neuroticism score at study enrollment vs. overall cognitive change over time. Negative values indicate a decline in global cognition. The red dashed line represents a linear regression model fit to the data; the shaded region represents the 95% confidence interval, indicating the precision of the estimated effect across the range of neuroticism scores. (B) A forest plot visualizing the relationship between neuroticism score vs. cognitive change within the core domains. Each point represents an odds ratio, with red points denoting statistical significance. The error bars indicate the 95% confidence interval. The associations between neuroticism vs. domain-specific cognitive change were quantified using linear regression modeling, adjusting for age at death, age at baseline, years of education, sex, and study.

Additionally, we examined the association between neuroticism and classic measures of AD pathology. While neuroticism demonstrated a nominal positive association with tau pathology (*p* = .04), it did not exhibit an association with Aβ pathology (*p* = .074) (**Supplementary Table S1**).

### Neuroticism is associated with gene expression changes in excitatory neurons

After establishing the association between neuroticism and AD in our ROSMAP sample, we leveraged the corresponding snRNA-seq data for downstream analysis. We performed a two-stage differential expression analysis to identify genes associated with neuroticism within neocortical cell types. In Stage 1, we examined gene expression in the seven major neocortical cell types and prioritized genes that were nominally (p <= 0.05) associated with neuroticism in our largest dataset (CUIMC1, n = 371) (**Table 2**). The prioritized genes from Stage 1 were advanced to Stage 2, in which we evaluated the reproducibility of these gene associations using a fixed-effects meta-analysis of the two remaining, independent datasets (CUIMC2, n = 172; MIT, n = 112).

**Table 2:** Two-stage analysis reveals two differentially expressed genes in excitatory neurons of individuals with elevated neuroticism scores.

| <u>Major Cell Type</u> | <u>Number of Discovery Genes</u><br>($p < 0.05$) | <u>Replicated Genes</u><br>($FDR < 0.05$) |
| --- | --- | --- |
| Astrocytes | 449 | 0 |
| Oligodendrocyte | 397 | 0 |
| OPCs | 693 | 0 |
| Inhibitory Neurons | 380 | 0 |
| Excitatory Neurons | 274 | **ADRA1B, LY6E-DT** |
| Microglia | 1859 | 0 |
| Endothelial Cells | 606 | 0 |

To explore whether the number of genes that are nominally associated with neuroticism differs across cell types, we examined the distribution of resulting p-values from the Stage 2 analysis. We found that excitatory neurons demonstrated a clear enrichment of genes with nominally significant associations with neuroticism, whereas the other neocortical cell types displayed near uniform p-value distributions (**Fig. 3A**). This relative enrichment suggests that excitatory neurons may be particularly susceptible to the effects of neuroticism.

**Figure 3:**
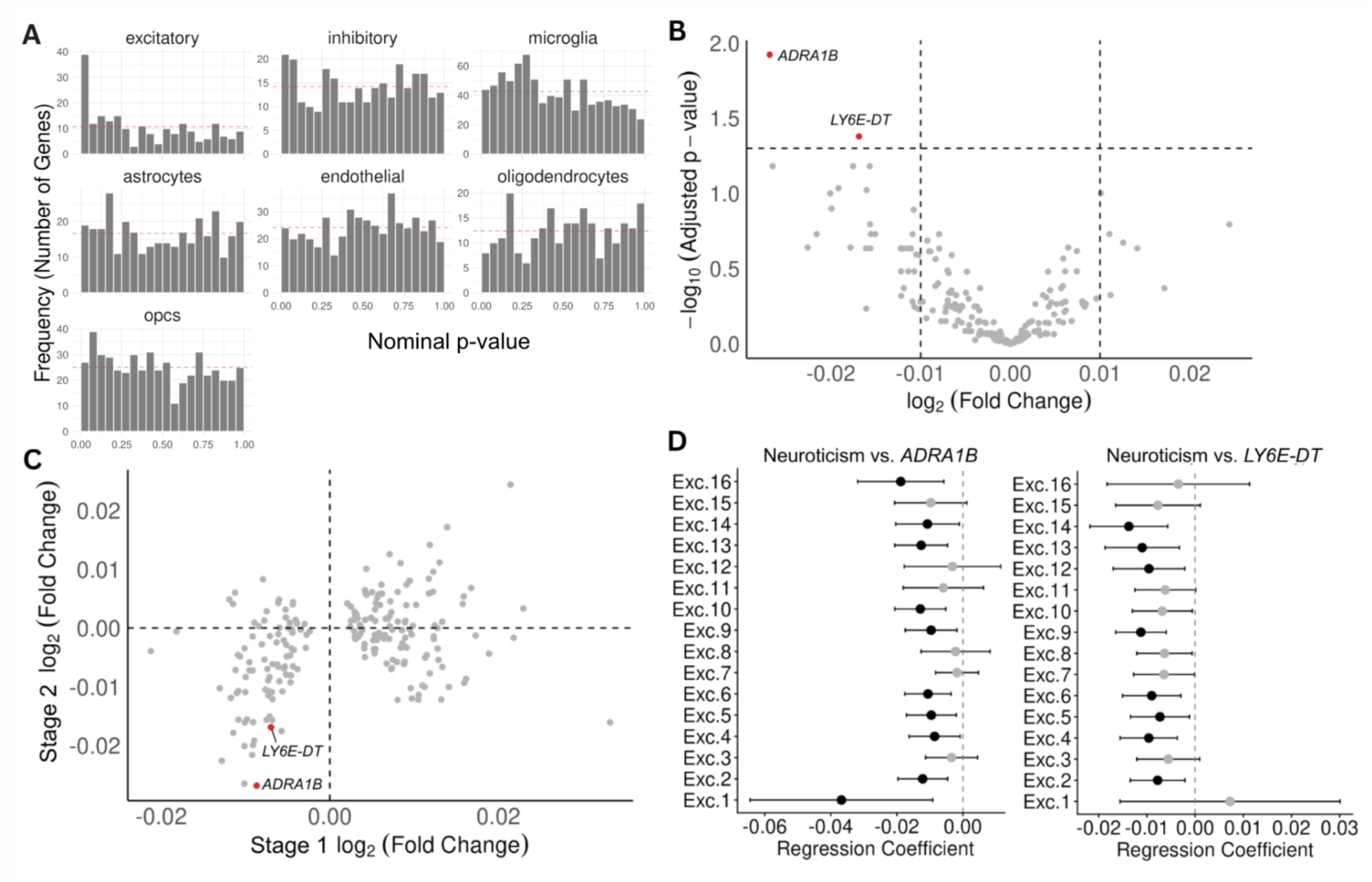
Neuroticism is associated with gene expression changes in excitatory neurons. (A) Histograms of the resulting nominal p-values from Stage 2 of the differential expression analysis, stratified by cell type. The horizontal red dashed line indicates the number of significant genes expected per cell type under the null hypothesis (α = 0.05). (B) A volcano plot visualizing differentially expressed genes in excitatory neurons, with significant genes denoted in red (FDR ≤ 0.05). The x-axis indicates the log_2_(Fold Change) of gene expression in individuals with elevated neuroticism scores. The y-axis indicates the -log_10_(Benjamini-Hochberg adjusted *p*-value) of the association between neuroticism score and gene expression. The vertical dashed lines are placed at log_2_(Fold Change) values of −0.01 and 0.01. The horizontal dashed line is placed at an adjusted p-value of 0.05 to indicate the significance threshold. (C) A scatter plot visualizing the directionality of differentially expressed genes in excitatory neurons in Stage 1 vs. Stage 2, with significant genes denoted in red (FDR ≤ 0.05). The x-axis indicates the log_2_(Fold Change) of gene expression from the Stage 1 output. The y-axis indicates the log_2_(Fold Change) of gene expression from the Stage 2 output. (D) For *ADRA1B* and *LY6E-DT*, the association between neuroticism score and gene expression was assessed at the cell subpopulation level. In the above forest plots, the x-axes show the regression coefficients for the neuroticism vs. gene expression associations. The points are colored by significance, with black points indicating significant associations (FDR ≤ 0.05).

After correcting for the testing of multiple hypotheses, the Stage 2 analysis revealed two differentially expressed genes, both of which were downregulated in excitatory neurons: *ADRA1B* (Benjamini-Hochberg adjusted *p* = 1.19 × 10^−2^) and *LY6E-DT* (Benjamini-Hochberg adjusted *p* = 4.17 × 10^−2^) (**Fig. 3B**). The directionality of these associations was consistent with the Stage 1 analysis (**Fig. 3C**). In a supplementary analysis, we explored whether these genes were associated with neuroticism within inhibitory neurons; however, we did not find evidence of an effect in this cell population (Benjamini-Hochberg adjusted *p* > 0.05).

To investigate whether a subset of excitatory neurons drives this overall effect, we examined the relationship between neuroticism and *ADRA1B* as well as *LY6E-DT* within each of sixteen predefined excitatory neuron subpopulations^10^. The association between neuroticism and *ADRA1B* was observed in ten of the sixteen excitatory neuron subpopulations, and the association between neuroticism and *LY6E-DT* was observed in eight (**Fig. 3D**). These findings suggest that the association between neuroticism and the expression of these genes is likely to be distributed across a range of excitatory neuron subpopulations.

### Neuroticism is linked to enrichment of a specific excitatory neuron subpopulation

Given the enrichment of neuroticism-associated gene expression changes in excitatory neurons, we then investigated whether neuroticism influences the composition of excitatory neurons at the subpopulation level. To address this, we examined the association between neuroticism and excitatory neuron subpopulation frequencies. This analysis revealed a specific excitatory neuron subpopulation with a significant positive association with neuroticism: Exc.12 (Benjamini-Hochberg adjusted *p* = 3.82 × 10^−2^) (**Fig. 4A, Supplementary Table S6**). These findings suggest that the Exc.12 neuronal subpopulation is more prevalent in the neocortex of participants with elevated neuroticism.

**Figure 4:**
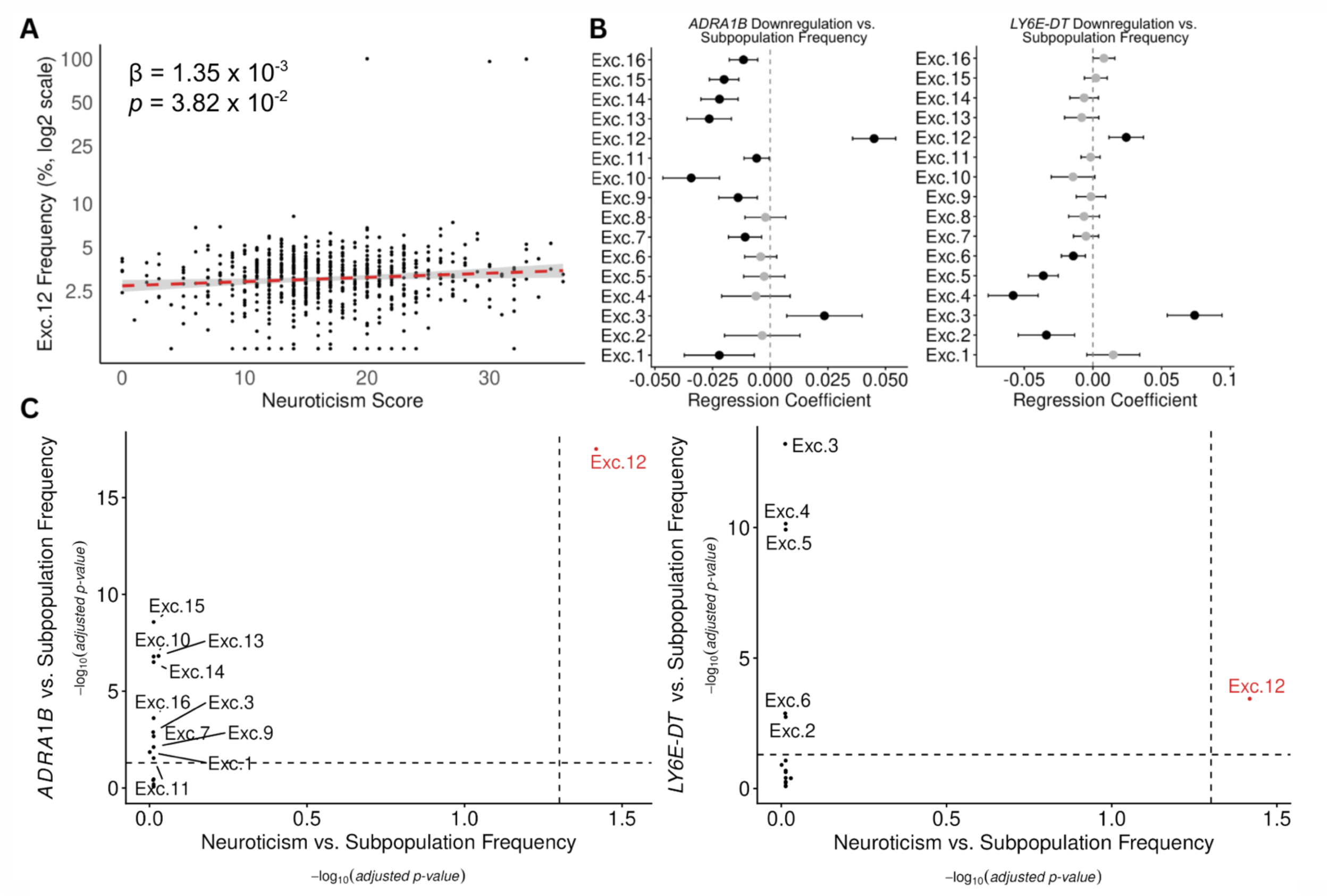
Neuroticism is linked to enrichment of a specific excitatory neuron subpopulation: Exc.12. (A) A scatter plot illustrating the association between neuroticism score and Exc.12 frequency. The x-axis indicates neuroticism score at baseline. The y-axis indicates Exc.12 frequency after log_2_ transformation. B) For each excitatory neuron subpopulation, the association between neuroticism-associated gene expression changes vs. subpopulation frequency was assessed using linear regression modeling, adjusting for demographic and technical confounders. In the above forest plots, the x-axes show the resultant regression coefficients from these analyses. The points are colored by significance, with black points indicating significant associations (FDR <= 0.05). (C) Each point represents an excitatory neuron subpopulation, with the significant subpopulation (Exc.12) indicated in red. The x-axis shows the -log_10_(Benjamini-hochberg adjusted *p*-value) of the association between neuroticism and subpopulation frequency, quantified using linear regression modeling. The y-axis shows the - log_10_(Benjamini-hochberg adjusted *p*-value) of the association between gene expression and subpopulation frequency, quantified using linear regression modeling. The vertical and horizontal dashed lines indicate the significance threshold (Benjamini-hochberg adjusted *p*-value = 0.05).

Further, we assessed whether the expression level of our two neuroticism-associated genes, *ADRA1B* and *LY6E-DT*, is associated with excitatory neuron subpopulation frequencies. We found that neuroticism-associated gene expression changes correlate with significant shifts in the composition of excitatory neurons (**Fig. 4B**), including an enrichment of the Exc.12 subpopulation (**Fig. 4C**).

To explore whether this neuroticism-associated subpopulation is also linked to neurodegeneration, we then tested the association between Exc.12 frequency and AD phenotypes, including a clinical diagnosis of AD, cognitive decline, Aβ pathology burden, and tau pathology burden. Exc.12 frequency was significantly associated with a clinical diagnosis of AD (*p* = 6.75 × 10^−3^) and cognitive decline (*p* = 5.16 × 10^−4^) (**Supplementary Table S7**). Furthermore, Exc.12 demonstrated a modest positive association with tau pathology burden (*p* = 1.72 × 10^−2^) and no association with Aβ pathology burden (*p* = 6.65 × 10^−1^). These findings were consistent with the associations observed between neuroticism and these AD phenotypes (**Supplementary Table S1**). Given that Exc.12 demonstrated evidence of enrichment in neuroticism as well as AD phenotypes, we selected this excitatory neuron subpopulation as a candidate for mediation analysis.

### The Exc.12 subpopulation mediates the association between neuroticism and AD, independent of Aβ and tau pathology

To assess whether the Exc.12 subpopulation mediates the relationship between neuroticism and AD, we performed a mediation analysis. We found that Exc.12 frequency demonstrates statistical evidence of mediation, accounting for 12.2% of the total effect of neuroticism on AD (*p* = 1.90 × 10^−2^) (**Fig. 5A**). Similarly, Exc.12 frequency was found to mediate the association between neuroticism and cognitive decline, accounting for 10.5% of the total effect (*p* = 2.09 × 10^−2^) (**Fig. 5B**).

**Figure 5:**
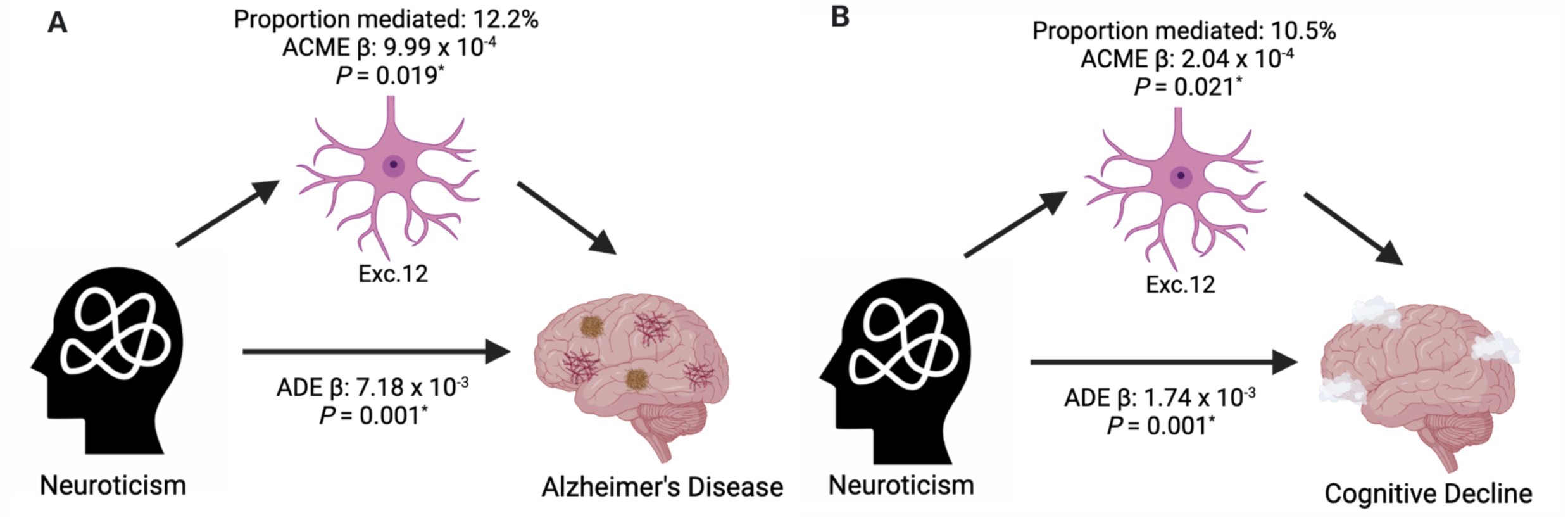
Exc.12 frequency mediates the relationship between neuroticism and AD, as well as the relationship between neuroticism and cognitive decline. (A) Causal mediation model illustrating the relationship between neuroticism and AD, mediated by Exc.12 frequency. The above diagram indicates the indirect effect of neuroticism on AD through Exc.12 (ACME), the direct effect of neuroticism on AD holding Exc.12 constant (ADE), as well as the percentage of the total effect that is mediated by Exc.12 (proportion mediated). (B) Causal mediation model illustrating the relationship between neuroticism and cognitive decline, mediated by Exc.12 frequency. The above diagram indicates the indirect effect of neuroticism on cognitive decline through Exc.12 (ACME), the direct effect of neuroticism on cognitive decline holding Exc.12 constant (ADE), as well as the percentage of the total effect that is mediated by Exc.12 (proportion mediated).

To investigate whether Exc.12 exerts this mediation effect through proteinopathy, we conducted additional mediation analyses of the relationship between neuroticism and tau pathology. However, these analyses revealed that Exc.12 does not mediate the relationship between neuroticism on tau deposition (*p* = 1.62 × 10^−1^) (**Supplementary Table S8**). These findings suggest that Exc.12 plays a role in mediating the association between neuroticism and AD independent of proteinopathic mechanisms.

## Discussion

Consistent with previous studies, we observed that neuroticism is associated with an increased risk of AD and accelerated cognitive decline^2–5^. Expanding on these findings, additional analyses revealed that the relationship between neuroticism and AD remains significant even after adjusting for Aβ and tau deposition in the brain. This robust association raised a question: what are the mechanisms through which neuroticism impacts the aging brain?

To address this question, we explored the molecular underpinnings of the relationship between neuroticism and AD using single nucleus transcriptomic data. We ran a two-stage differential expression analysis: taking an exploratory approach in Stage 1, followed by validation in Stage 2. This analysis revealed that excitatory neurons are enriched for neuroticism-associated genes and may be implicated in the genetic manifestations of this personality trait.

After correcting for the testing of multiple hypotheses, we identified two genes associated with neuroticism, both of which were downregulated in excitatory neurons: *ADRA1B* and *LY6E-DT*. *LY6E-DT* is a long non-coding RNA whose function is underexplored and has not been previously implicated in neurological disease. *ADRA1B* encodes the α1β-adrenergic receptor^21^, a component of the noradrenergic signaling pathway involved in stress response, alertness, and attention. Altered expression of *ADRA1B* may relate to altered noradrenergic signaling secondary to chronic stress exposure in individuals with neuroticism. A study investigating the role of noradrenergic dysfunction in neurodegeneration found that *ADRA1B* knockout mice displayed deficits in memory consolidation, linking this gene mechanistically to memory circuits that may contribute to dementia^22^.

Given that neuroticism demonstrated differential gene expression in excitatory neurons, we investigated the impact of neuroticism on this cell type further. Our analysis revealed a subpopulation of excitatory neurons that is more prevalent in individuals with elevated neuroticism: Exc.12. The Exc.12 subpopulation frequency is also linked to neuroticism-associated gene expression changes in *ADRA1B* and *LY6E-DT*, offering support for a transcriptomic link between the personality trait and this cell subpopulation. Further analyses revealed that Exc.12 mediates 12.2% of the effect of neuroticism on AD through non-proteinopathic mechanisms.

A cell atlas of the aged prefrontal cortex helps contextualize the function of the Exc.12 subpopulation^10^. The atlas annotations reveal that Exc.12 expresses the marker gene *FEZF2* and localizes to cortical layers 5 and 6. These layers are rich in subcortical projecting neurons, which play a role in communicating between the cortex and structures such as the amygdala, thalamus, and hippocampus^23^. The localization of Exc.12 to the deep cortical layers suggests that this subpopulation may play a role in top-down communication to subcortical structures. Furthermore, the top differentially expressed pathways in Exc.12 relate to synaptic organization, density, and signaling^10^. These pathways are significantly downregulated in the Exc.12 subpopulation, suggesting that this subpopulation may be associated with impaired synaptic integrity.

Since these analyses were conducted using cross-sectional data collected at autopsy, we cannot comment on causality; however, our statistically rigorous mediation analyses are helpful in generating hypotheses. We hypothesize that neuroticism may lead to an increased frequency of the Exc.12 subpopulation in older age, contributing to an increased risk of AD and accelerated cognitive decline. Since neuroticism does not change meaningfully after 25 years of age, it is reasonable to propose that it precedes cognitive decline later in life^1^. We posit that the relative frequency of the Exc.12 subpopulation increases over the course of life secondary to chronic stress exposure in individuals with elevated neuroticism. Therefore, the Exc.12 subpopulation may represent a dysfunctional neuronal state. However, given that our data are cross-sectional, we cannot exclude the possibility that the increased frequency of Exc.12 is a feature of the neural circuits that predispose people to neuroticism and thus may have emerged earlier in life, such as during nervous system development.

Validation and further exploration of these findings in cell culture or mouse model systems could guide therapeutic development and empower targeted interventions. For example, gaining a better understanding of the role and function of Exc.12 could inform the development of drugs that protect or even restore synaptic plasticity in the aging brain. Furthermore, these findings could aid in targeting existing drugs to patients who are most likely to reap clinical benefit. Consider donepezil, a commonly prescribed medication in the symptomatic management of AD. Donepezil is a cholinesterase inhibitor and acts directly at the synapse to prevent breakdown of acetylcholine. Though donepezil has been shown to improve cognition at the population level, individual patient responses vary widely and range from marked to minimal improvement^24^. Additionally, side effects of this medication impact quality of life and can include nausea, vomiting, syncope, and dizziness^25^. Understanding how neuroticism impacts synaptic health could help identify patients who are more likely to benefit from initiation of symptomatic treatments such as donepezil.

While this work offers important insights into the role of neuroticism in neurodegeneration, there are limitations to consider. First, the snRNA-seq data used in our analyses are cross-sectional and thus prevent us from resolving causal relationships. Second, the data were derived from a predominantly female cohort of European ancestry, which limits our ability to generalize these findings to a broader, more diverse patient population. Third, all sequencing data were obtained from the dorsolateral prefrontal cortex, and therefore our findings are specific to this brain region; they may not capture transcriptional changes in other brain regions involved in neuroticism.

Despite these limitations, there are key strengths of this study as well. The large sample size of the study allowed for sufficient power to detect subtle effects and pursue rigorous mediation analyses. Additionally, the three separate sequencing cohorts were well-suited for a multi-stage analysis, allowing for validation of genes of interest. Furthermore, the single-nucleus resolution of our transcriptomic data enabled us to identify differences in the impact of neuroticism on the various neuronal and glial cell types.

In conclusion, this study sheds light on the mechanisms through which neuroticism impacts the aging brain. We demonstrate that neuroticism is associated with a higher frequency of the Exc.12 subpopulation, which mediates a meaningful portion of the trait’s total effect on cognition and risk of AD. Importantly, Exc.12 exerts this mediation effect independent of Aβ and tau accumulation, indicating that neuroticism may contribute to disease risk through mechanisms that extend beyond classic proteinopathy. While experimental models are needed to validate these findings, our results support a cellular link between neuroticism and neurodegeneration, highlighting a specific excitatory neuron subpopulation as a compelling candidate for further investigation.

## Consent Statement

Participants signed an informed consent, a repository consent, and an Anatomical Gift Act to donate their brains after death. Both studies were approved by an Institutional Review Board at Rush University Medical Center.

## Funding Statement & Acknowledgements

We thank the ROSMAP participants and their families for their invaluable contribution to this research. The ROSMAP effort is supported by the following grants: P30AG10161, P30AG72975, R01AG17917, R01 AG015819, U01 AG072572, and U01 AG046152.

